# Determination and correction of aberrations in full field OCT using phase gradient autofocus by maximizing the likelihood function

**DOI:** 10.1101/2020.05.12.089128

**Authors:** Vasily Matkivsky, Alexander Moiseev, Pavel Shilyagin, Alexander Rodionov, Hendrik Spahr, Clara Pfäffle, Gereon Hüttmann, Dierck Hillmann, Grigory Gelikonov

## Abstract

A method for numerical estimation and correction of aberrations of the eye in fundus imaging with optical coherence tomography (OCT) is presented. Aberrations are determined statistically by using the estimate based on likelihood function maximization. The method can be considered as an extension of the phase gradient autofocusing algorithm in synthetic aperture radar imaging to 2D optical aberrations correction. The efficiency of the proposed method has been demonstrated in OCT fundus imaging with 6λ aberrations. After correction, single photoreceptors were resolved. It is also shown that wavefront distortions with high spatial frequencies can be determined and corrected.

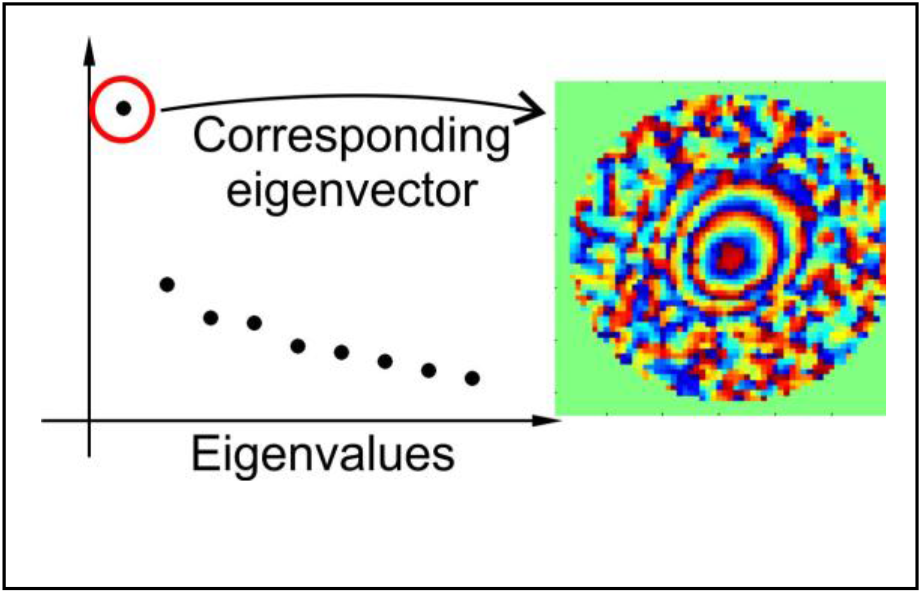

**Graphical Abstract for Table of Contents:** **[Text.** This work is dedicated to development a method for numerical estimation and correction of aberrations of the eye in fundus imaging with OCT. Aberration evaluation is performed statistically by using estimate based on likelihood function maximization. The efficiency of the proposed method has been demonstrated in OCT fundus imaging with 6λ aberrations. It has been shown that spatial high-frequency wavefront distortions can be determined]

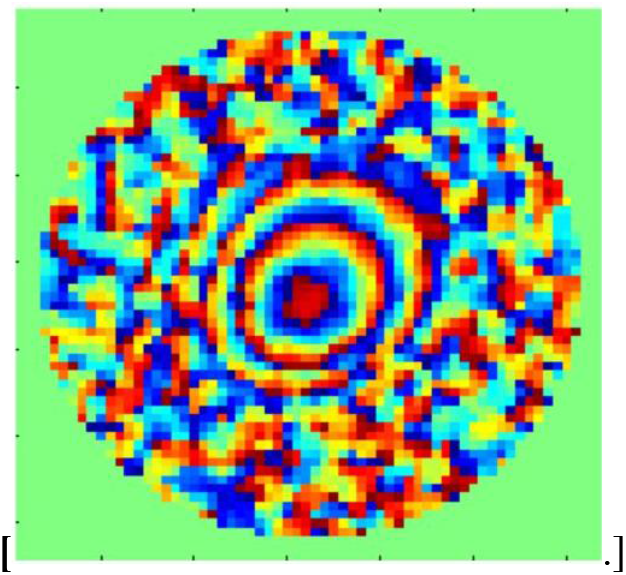

## 1 INTRODUCTION

One of the challenges in ophthalmoscopy is high-resolution imaging of the human retina, which is hindered by wavefront distortions (optical aberrations) introduced by the eye into the optical imaging system. To determine and compensate for the influence of these aberrations, adaptive optics (AO) is being successfully used [1]. This technique requires additional elements in the optical setup: means for projecting a guide-star onto the retina, a wavefront sensor, and a wavefront corrector. With the advent of imaging techniques which record scattered amplitude and phase of the electrical field, it has become possible to replace AO elements by numerical image restoration algorithms for determining and correcting optical aberrations numerically. Several ways for aberration correction without using additional optical components have been demonstrated. In digital holography [2], the hologram of a guide-star was used for aberration detection and correction [3,4]. In [5,6] aberrations were detected numerically by image quality metric optimization and were corrected by using a deformable mirror. When amplitude and phase are detected without object motion, both detection and correction of aberration can be done numerically. In [7–9] wavefront distortions were considered as a sum of Zernike polynomials. Zernike decomposition coefficients are obtained using iterative optimization of an image quality metrics. The drawbacks of this method are a high computational cost and the risk of ending in a local minimum of the image quality metrics, which may result in poor image quality. Also the combination of hardware AO and optimization method can be effective [10]. Hardware AO allows to collect more photons and numerical correction can easily address high-order aberrations.

Recently, several methods for all numerical aberration correction have been proposed that do not require solving an optimization problem. One of the approaches is to use different sub-apertures to reconstruct wavefront distortions [11,12]. This technique suffers from a trade-off between precision and resolution and has difficulties determining higher-order aberrations of the eye. Using multiple randomly choosen chosen subabertures [13] or a two stage procedure improved the results [14,15]. In the latter, the first stage determines the aberrations by the sub-aperture correlation technique, and the second stage selects single photoreceptors within the image and uses them as a guide-star for final aberration detection. This allowed determining distortions of 24 Zernike terms wavefront approximation with RMS error of about 0.54 *λ* for 3.5 mm pupil diameter in the work [15] and a RMS of about 0.48 λ for 14 Zernike terms and 7 mm pupil diameter in the work [14].

Here, we propose the alternative method. We seek not for a single scatterer but for multiple point-like scatterers which need not necessarily be isolated as guide stars but may overlap with other imaged structures. Based on images of these scatterers, using the maximum likelihood method, the optical aberrations are estimated, and a corrected image is reconstructed. This method is analogous to the phase gradient autofocusing (PGA) used in synthetic aperture radar (SAR) imaging [16], but here it is applied to 2D phase errors. This method enables estimation of large eye aberrations with RMS of about 0.85 λ (1.15 λ with defocus) and 42 Zernike terms. Extraction of high-frequency aberrations up to 117 Zernike terms are demonstrated on a model object.

Initially, this approach was applied to OCT data for digital refocusing [17]. In our earlier work, we used a similar approach for compensating the influence of aberrations in digital holography, motion-compensation in full-field swept-source OCT, and dispersion in spectral-domain OCT imaging [18–20]. However, the direct application of this approach to ophthalmic data gave poor result. Below we demonstrate how this method may be refined to the level that will allow determining strong eye aberrations. The proposed method is compared with optimizing image quality metrics.

## 2 Materials and Methods

### 2.1 Data acquisition

Data generated with two setups were used to demonstrate the algorithm. A digital holography setup allowed introducing particular high-order aberrations and was used to compare the efficacy of the improvement algorithm for detecting wavefront distortions and compare it to results from our previous approach [18]. The setup had an off-axis Michelson interferometer with a semiconductor laser source at a wavelength of 847 nm (fig. 1). Aberrations were introduced by a transparent mask with exactly known refractive index and geometry [21], which introduces the wavefront distortions presented in fig. 1b. The distortion has a maximum amplitude of approximately 15 rad.

**FIGURE 1.**
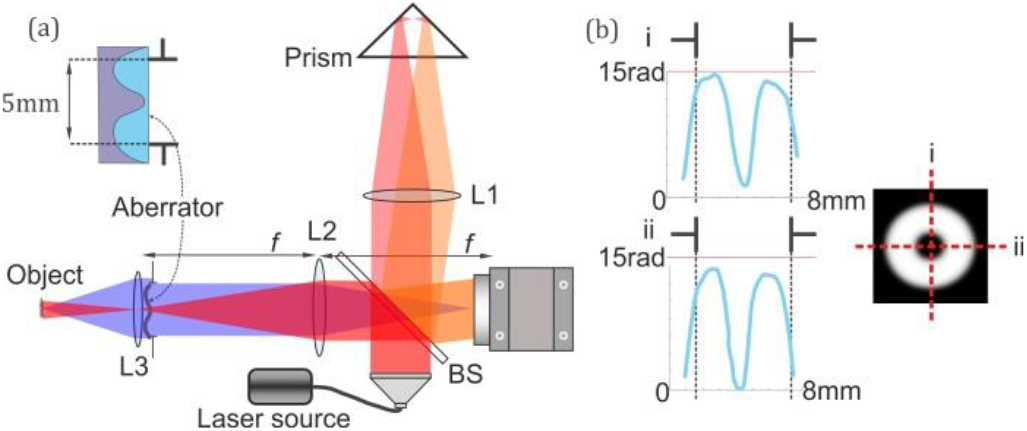
Digital holography setup. a) Light from semiconductor source with wavelength 847 nm (Superlum BS 840) is split into reference and sample arm. The negative USAF1951 test target (Object) is illuminated through the lens system L2 (Thorlabs AC508-075) – L3 (Thorlabs AC127-030). The white composition was applied to the objects’ surfaces for the additional scattering. Backscattered light (blue) is focused onto the camera (Thorlabs DCC 1240M) by the optical system comprising the lenses L3, L2 and the aberrator placed in the focal plane of the lens L2. The camera is placed in the back focal plane of the lens L2. b) Wavefront distortion profile, induced by the aberrator.

To demonstrate the efficacy of the proposed method for aberration correction in a real application, data from fully phase-stable retinal imaging by full field swept source OCT have been used [8].

### 2.2 Aberration detection and correction

Wavefront distortions are caused by the radiation passage through L3 with the aberrator or the anterior section of the human eye both of which are introduced by the phase error *ϑ*(*k_x_, k_y_*) in the Fourier domain of the optical imaging system. A convolution with the Fourier transform of 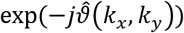 gives a corrected image, where 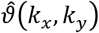 is the estimate of the true phase-error function *ϑ*(*k_x_, k_y_*) and (*k_x_, k_y_*) are discrete spatial frequencies. An imaging model is needed to find the estimate of 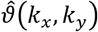.

The aberration-free field *E*_0_(*x, y*) of scattered light can be represented as a sum of diffraction limited point spread functions (PSF) *A*_0_(*x, y*):

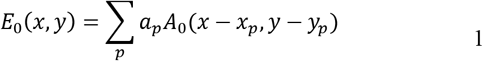

where (*x_p_, y_p_*) is the coordinate of the PSF’s center, and *a_p_* is its complex amplitude. With denoting the Fourier transform, the aberrated image in the frequency domain can be represented as:

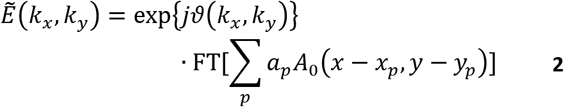

Thus, the image field of the complex amplitudes can be written in the form:

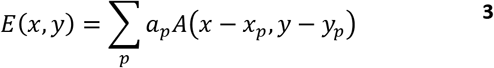

In contrast to eq. (1), *A*(*x* − *x_p_, y* − *y_p_*) is an aberrated PSF. Note that the recorded image consists of several overlaid and distorted PSFs (images of a point-like scatterers), each of which contains information about the phase error *ϑ*(*k_x_,k_y_*). We will assume point-like scatter as one or several individual scatterers separated by distance less than diffracted limited PSF size. On the one hand, images of scatterers overlap due to aberrations or the limited resolving power of the optical system, making it impossible to extract an isolated bright scatterer image. On the other hand, the multiplicity of these images generates redundant information about the PSF. The proposed algorithm relies on the presence in the image of some dominant scatterers, which is the case in the practically important case of the retina photoreceptors. In a more general case one can hope that the image will comprise of some contrast structures oriented in different directions which is the case in USAF1951 target, successfully restored with the proposed algorithm.

To make use of this information the maximum likelihood estimation method is employed, similar to the image reconstruction in SAR imaging [16]. We will divide the procedure into several steps for clarity. The algorithm of this procedure is shown schematically in fig. 2.

**FIGURE 2.**
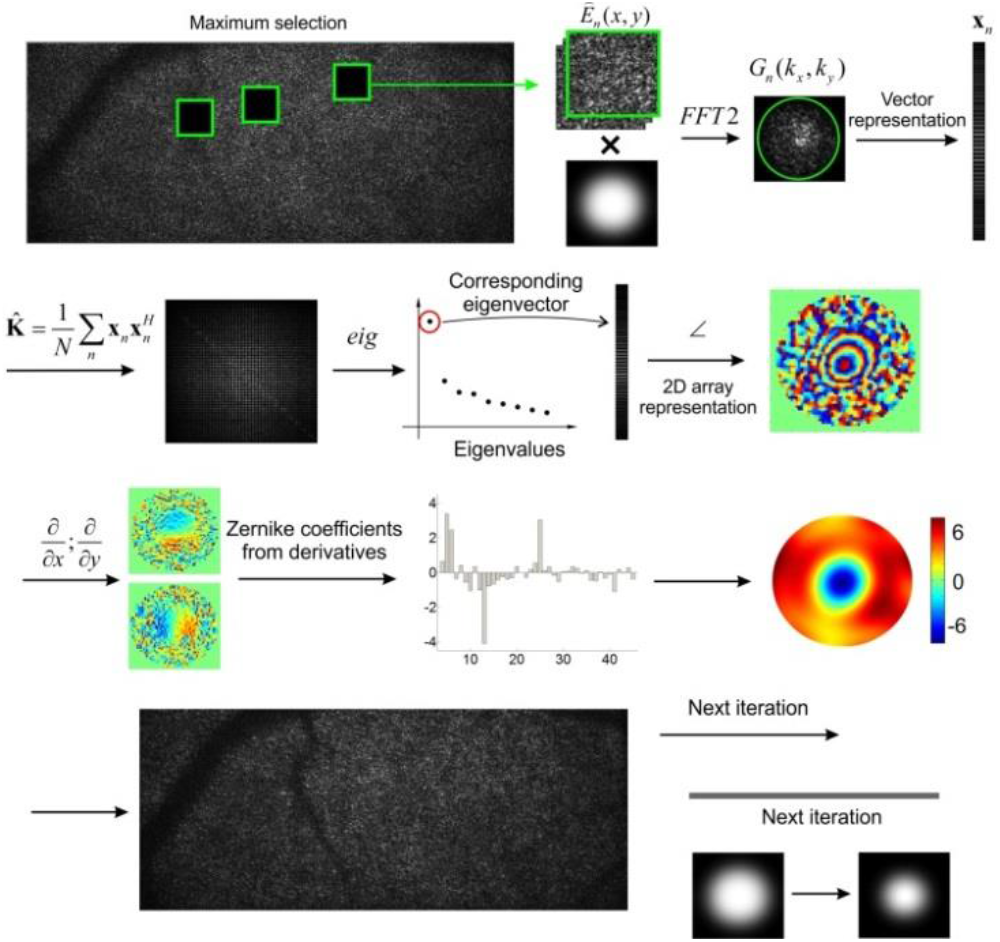
Data flow of the PGA with eigenvector estimator.

Step 1. Sub-images are cut around the points of local maximum brightness. Several local maxima (typically about a hundred) of the absolute values of the field *E*(*x, y*) should be found in the image plane. The maxima are found iteratively starting with the point with maximal field intensity. The next maximum is found as the point with maximal field intensity which lies not closer than 2R (R is radius of the function, see step 2) distance from any of the previously found maxima. The search continues iteratively until the field value in the next maximum will be less than the maximal field value in the image divided by 2.5 or until the total number of found maxima will exceed 250. Overall, a larger number of selected sub-images with high Signal-to-Noise ratio (SNR) will lead to the better aberation estimation, at the cost of increased computational time. At the same time selection of local maxima with low SNR will negatively affect the algorithm performance. The stopping criteria in the present study were selected empirically as a trade-off between computational time and algorithm performance. The neighborhood of 64×64 pixels of each maximum is placed in a separate array. This gives a set of sub-image fields 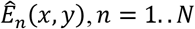.

Step 2. Next, 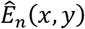 is multiplied by an apodization window function win(*x, y*) and Fourier-transformed:

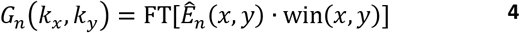

The apodization function win(*x, y*) is defined by:

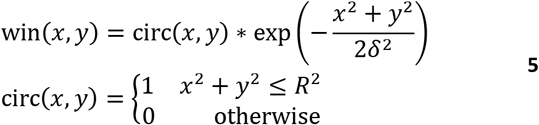

*R* is the radius of the circ function, which should contain main parts of the aberrated PSF, and * denotes the two-dimensional convolution. The radius will decrease during the following iterative reconstruction, as the image becomes sharper. The area of the subimages defined in step 1 is chosen such that the distance between adjacent sub-images will be no less than 2*R*. One should note that decreasing the *R* value leads to the increasing of the potential subimages used for the aberration estimation. The initial value of *R*, its decrease with each iteration, and *δ*^2^ depend on the experimental conditions (severity of aberrations, pixel size and optical magnification) and are chosen empirically. The initial radius and decreasing step were 46 and 2 pixels respectively for the USAF target and 44 and 4 pixels for the retina. The convolution with a Gaussian function is performed to reduce the diffraction effect of the sharp boundary of the circ function and *δ*^2^ sets the border smoothness. On the basis of the set of Fourier transforms *G_n_*(*k_x_,k_y_*) of the sub-images, it is necessary to calculate the covariance matrix for determining the aberrations. Next, we show that maximizing the likelihood function problem mathematically reduces to finding the first eigenvector of the covariance matrix. For its calculation we represent *G_n_*(*k_x_,k_y_*) as a column vector **x**_*n*_ = [*g*_*n*1_, *g*_*n*2_,…, *g_nS_*], where s enumerates over wavenumbers k_x_,k_y_ inside the aperture of G_n_:

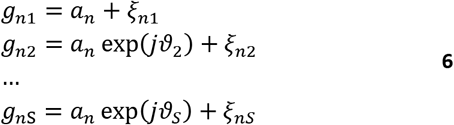

Here, *g_ns_* = *G_n_*(*k_x_,k_y_*), *a_n_* is a complex constant, and *ϑ*_1_ is assumed to be zero. The term *a_n_* exp(*jϑ_S_*) corresponds to the Fourier transform of the aberrated PSF. The term *ξ_ns_* is related to the sum of different parts of closest scatterers in the cut-out area. The behavior of *ξ_ns_* can be considered as noise, because different parts of many point-like scatterers with different amplitudes fall in the cutout area (sub-image), so that according to the central limit theorem the real and imaginary parts can be considered to be Gaussian distributed and the noise should follow speckle statistics. Each column vector **x**_*n*_ is statistically independent due to non-overlapping sub-images. The expression (6) coincides with the statistical model from the reconstruction of in synthetic-aperture-radar images [16] and a maximum likelihood (ML) estimator can be obtained analogously:

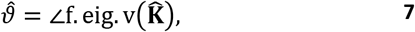

where ∟ denotes the argument of a complex value, f. ei g. v is the first eigenvector, i.e., the eigenvector corresponding to the largest eigenvalue, and 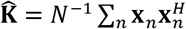 is the sample covariance matrix. This result can be interpreted as follows. The components of the covariance matrix, using eq. (6), can be found as:

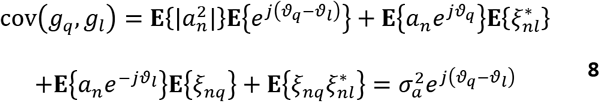

Here, **E** denotes the expectation value and 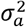 is the variance of *a_n_*, *q,l* ∈ 1..*S*. It can be seen that the 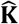 components contain various phase differences plus other terms which must tend to zero as *N* → ∞. The first eigenvector correlates with the most significant structure in the matrix, which corresponds to the wavefront distortion. This can be interpreted as the principal component analysis, where the first component contains the aberration phase and the other components contain noise. A simplified estimate for the wavefront distortions function using only adjacent pixels can also be obtained (see [16]):

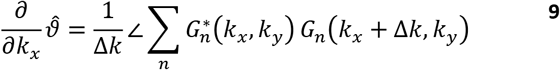

There Δ*k* = 1 for the utilized discretization. Differentiation by *k_y_* is omitted for brevity.. This formula allows us to obtain estimates of distorted phase gradients without calculating the covariance matrix and eigenvectors. Using the weighted least-squares technique [22] we can find the optical aberrations phase without any polynomial fitting. One should note that this technique was used for the phase calculation from the gradients estimated with the PGA. For short, we will call the estimation based on formula (7) an eigenvector estimator and formula (9) an adjacent pixel estimator.

The use of the covariance matrix allowed us to extract more information about the wavefront distortions at each iteration, which significantly increased the speed of the algorithm convergence and the resolution of the final image (see the section Results).

To increase the speed of the algorithm, the original covariance matrix was split into several smaller matrices by splitting the image aperture as shown in fig. 3. The most computationally expensive operations in the algorithm are: 1) obtaining a sample covariance matrix 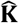 (calculation and summation of *N* products of 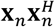) and 2) calculation of the eigenvectors of these matrices. For the first operation, dividing the aperture into *M* equal subapertures reduces the number of operations by *M* times. The speed of eigenvectors calculation also increases (by more than *M* times in practice). In addition, there are two positive features of these operations splitting the image aperture: 1) the possibility to parallelize the two operations described above and 2) an improvement of the speed of algorithm convergence and of the final image quality. We suppose that this is related to phase decorrelation between the remote parts of the field in the aperture.

**FIGURE 3.**
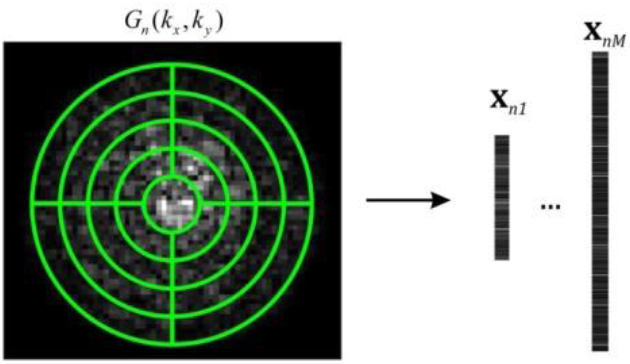
Scheme of image aperture splitting.

Step 3. The estimate 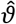 is differentiated along the *x* and *y* directions:

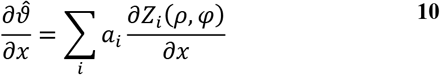

Differentiation by *y* is omitted for brevity. In the matrix form it can be represented as:

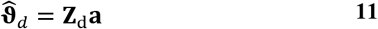

where 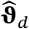 is the vector of derivatives of the estimate, **Z**_d_ is the matrix of the Zernike polynomials derivatives and **a** is the vector of Zernike coefficients. Next, **Z**_d_ can be represented as **Z**_d_ = **UDV**^T^ using singular value decomposition, and the coefficients vector **a** can be found as [23]:

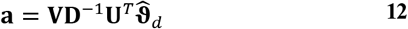

This avoids unwrapping in the phase matching procedure and gives a smoothed estimate. With the use of the Zernike coefficients, the estimate is extrapolated to the full grid (at Step 1 the covariance matrix and the estimate 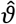 were found for the 64×64 grid).

At the first iteration, the corrected complex field *E*^1^ can be obtained in the form (index^1^ is the iteration number):

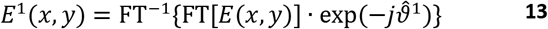

Next, the window is decreased in size and the procedure started anew. For a smaller window size, the signal-to-noise ratio (SNR) will increase and the estimate 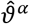 (where *α* is the iteration number) can be found more accurately. We used 8 iterations with a fixed decrease in the window size for each iteration. With a larger number of iterations (with a smaller step) the result is slightly better. The number of iterations was empirically chosen and set as one of the algorithm parameters.

One of the major problems is to determine the true centers of strong point scatterers in strongly aberrated images. For this, the image aperture is decreased numerically to a half of the initial radius in the Fourier domain and defocusing of the obtained image is compensated by the optimization method. Using this method in the considered case is justified, since it is much faster to determine one Zernike coefficient (not necessarily exactly) than several dozen of coefficients. This procedure improves the performance of the PGA method. Then, the corrected image with a smaller aperture is obtained by the PGA method. This small aperture image is used to find the coordinates of local maxima. The found maxima coordinates are used in the compensation algorithm with the full aperture image. Without these manipulations, the compensation algorithm gives poor results. Variations of the aberrations over the field of view do not significantly influence the compensation procedure, presumably because of the relatively small field of view.

The proposed algorithm was compared with the image quality optimization method using Shannon entropy of the central part of the restored images 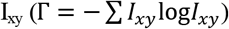. Its size was 512×512 in the case of the USAF target and 256×256 in the case of retina imaging. Smaller window size led to a significant deterioration in the convergence of the algorithm in both cases. The search for the coefficients in optimization method was performed with the Quasi-Newton method realized in MATLAB function fminunc. Gradients were calculated using method from [24] which allows to increase both computational speed and algorithm performance:

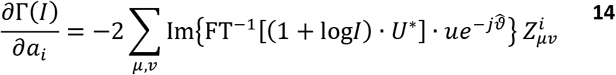

Where 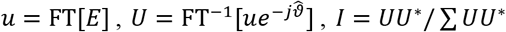. In case of the eye aberation correction aperture reduction technique was applied.

## 3 RESULTS AND DISCUSSION

The results of detecting the aberrations induced by the aberrator at numerical aperture (NA) of 0.08 are presented in fig. 4. The aberrations have a relatively high frequency: the Zernike polynomials up to the 14th radial degree, i.e., 117 degrees of freedom, were used to fit them.

**FIGURE 4.**
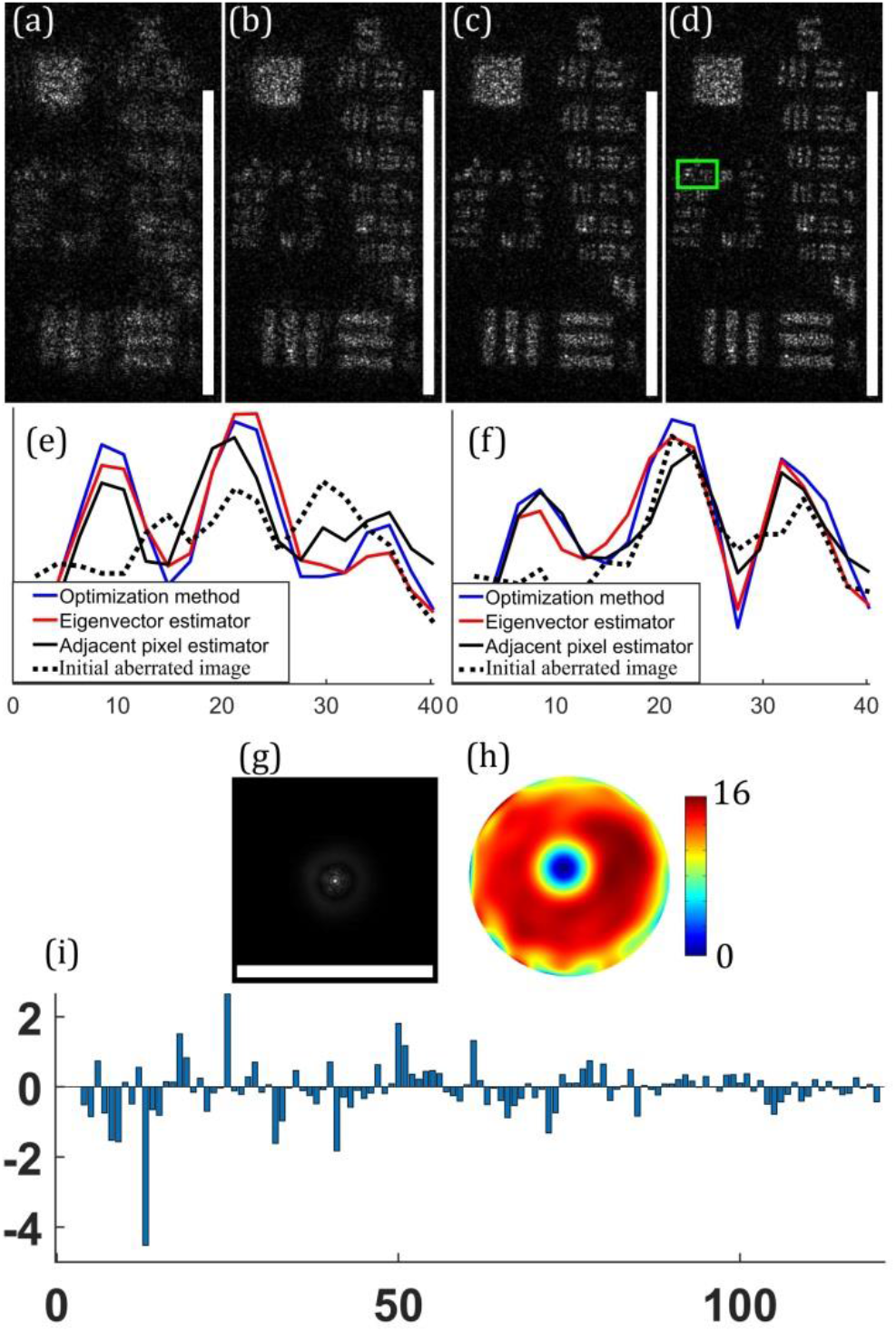
Digital holography image of the USAF 1951 before and after aberration removal. a) Before aberration correction, b-d) after aberration correction: using adjacent pixel estimator (b), using eigenvector estimator (c), using optimization method (d). Group 6 image averaging (in the green box) along: e) horizontal, f) vertical direction, g) point spread function, corresponding to detected wavefront distortion, i) Zernike coefficients. Scale bars are 0.5 mm

The proposed method was compared with optimization method [8] and adjacent pixel estimator [18]. It is interesting to compare the effectiveness of the optimization and the proposed methods in this case, in addition, it is interesting to compare the results with those of the previously performed work using the adjacent pixel estimator [18]. It can be seen that the here proposed PGA with eigenvector estimator (Figure 4c) and the optimization method (Figure 4d) provide close image quality, while PGA with adjacent pixel estimator (Figure 4b) works markedly worse. The compensation procedure took 1.6, 2, and 8.3 seconds for the PGA with adjacent pixel estimator, PGA with eigenvector estimator and optimization methods respectively, in the cases shown in fig. 4 (b), (c), (d), respectively (Intel Core i7 8700K platform, Matlab development environment). In Figure 5 one can see the detailed comparison of the proposed PGA with eigenvector estimator and optimization method for the various number of utilized Zernike polynomials. One can see that while the optimization method outperforms the proposed algorithm in terms of Shannon entropy, the proposed method requires less computational time. Visual analysis of the results in the same time shows comparable resulting image quality for the two methods. Compared to optimization method, the PGA works a little worse, but faster. In addition, execution time of optimization method has big variance, while PGA method has approximately the same execution time.

**FIGURE 5.**
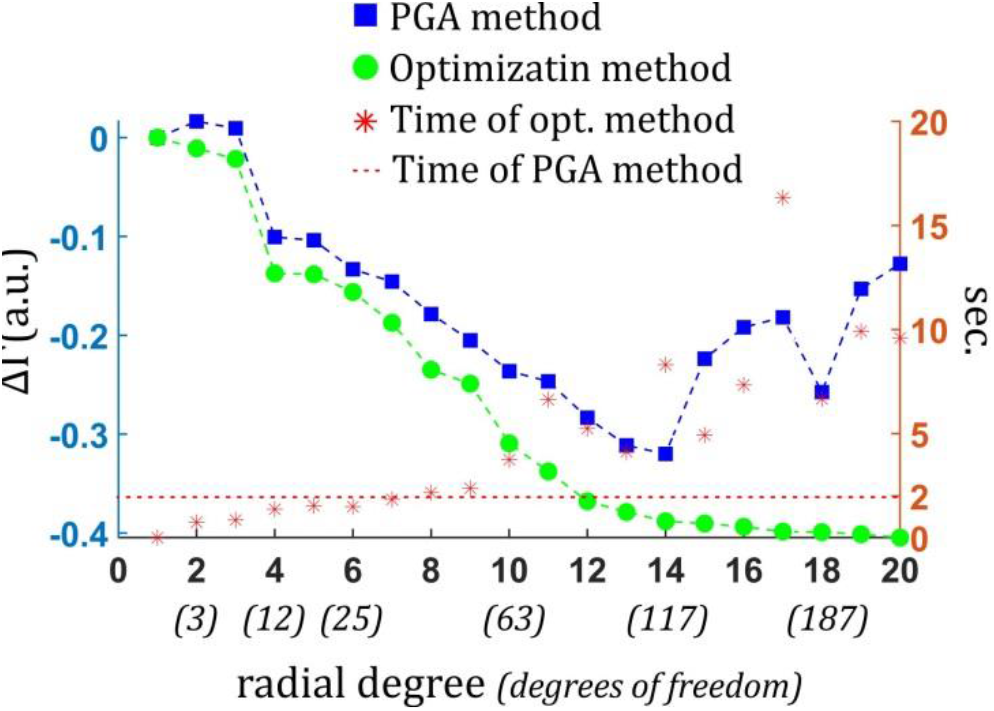
Performance comparison of PGA and optimization methods using different radial degrees of Zernike polynomials. Difference in Shannon entropy between the restored and initial images (left) and computational time (right) for the various radial degrees of Zernike polynomials with corresponding number of the calculated coefficients (in parenthesis) (x-axis). Blue line with square markers represents Shannon entropy for the PGA method, green line with circle markers represents Shannon entropy for the optimization method, red asterisks represent computational time for the optimization method, red dashed line represent computational time for the PGA method.

From Figure 5 one can see that the most significant improvement in terms of Shannon entropy corresponds to the usage of the first 12 polynomials, which include such aberrations as defocus, spherical aberration, coma and astigmatism. At the same time further increase of the polynomial degree improves both Shannon entropy and image quality. Since for the proposed PGA with eigenvector estimator such an increase in Zernike polynomial degree does not lead to the increased computational time, 117 degree polynomials were used in the present study. One can also see, that while for the optimization method higher degree of utilized polynomials leads to the further improvement of the Shannon entropy, for the proposed PGA method utilizing Zernike polynomials with radial degree higher than 14 leads to the degradation of the results, which may be attributed to the increased influence of the noise on the high-frequency components of the estimate.

The code was partially optimized but no parallelization was performed. It is worth noting that the optimization method was implemented using standard Matlab functions, which provides relatively good performance. The PGA method is relatively complicated for high-performance programming as it comprises several stages and operations with data. We believe that using a programming language such as C with good optimization will greatly improve efficiency, while the performance of the optimization method will not increase much.

The results of the aberration detection and correction for the data of *in vivo* human retina imaged with an NA of 0.2 [8] are presented in fig. 6. The compensation procedure for the image with a large NA aperture takes 5 seconds. The increase in computational time in comparison with the low-aperture case is caused by the increasing size of the covariance matrix. The computational time for the optimization method at the same time decreased to the 5 seconds, which was caused by the possibility of utilizing smaller window size (256×256) and smaller number of estimated parameters in comparison with the low-aperture case

**FIGURE 6.**
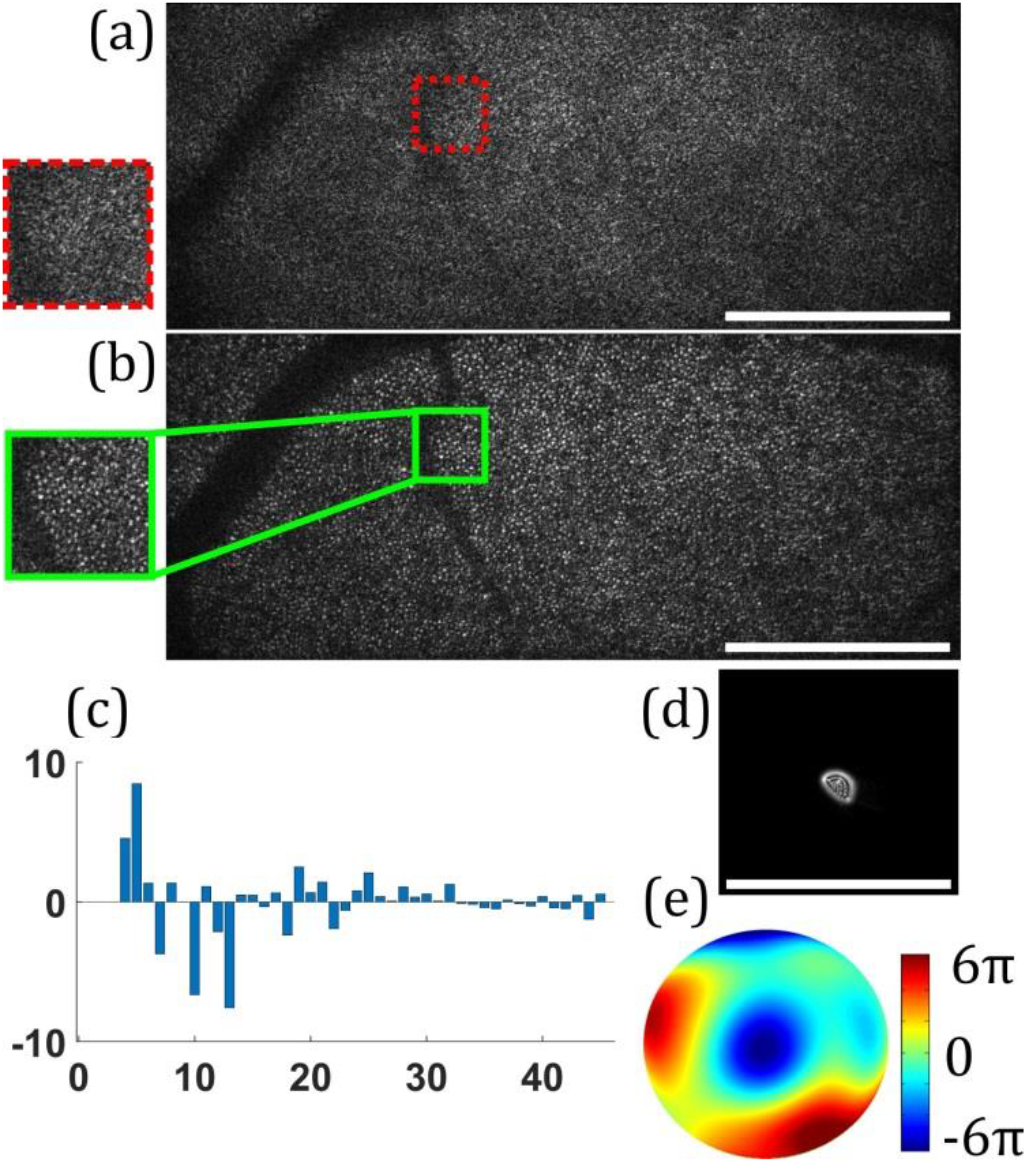
Retina image acquired by FF-SS-OCT at NA 0.2 after aberration removal. a) Image before aberration correction and b) image after aberration correction. The insets show magnifications of a small area. c) Zernike coefficients, d) calculated point spread function and e) wavefront distortion. Scale bars are 0.5 mm

After aberration correction the random speckle pattern is converted to the regular photoreceptor arrangement. The Yellott’s rings [25] in the Fourier transform of the reconstructed images and their averaging around the image centres (on a logarithmic scale) are presented in fig. 7. These rings are caused by the dominating spatial frequencies of the photoreceptor arrangement. The higher their amplitude, the sharper the reconstruction. It can be seen that the use of the eigenvector estimator increases the SNR by an average of 2.1 dB (averaging was performed between the vertical dashed lines in fig. 7e) compared to the adjacent pixel estimator and by 4 dB compared to the uncorrected image. The image obtained by the optimization method shown in fig. 7d is a little better that the image 7c, and the SNR is increased by 0.32 dB (the corresponding line in fig. 7e is absent due to the fact that it will be very close to the red line). It is worth noting that high-frequency polynomials play quite a significant role in our case. In figure 8c Shannon entropy dependence Zernike polynomials are presented for both PGA with eigenvector estimator (blue line with square markers) and optimization method (green line with circle markers). In this case one can see that the PGA with eigenvector estimator outperforms the optimization method for the majority number of utilized polynomials. The biggest improvement in Shannon entropy corresponds for the radial degree of 6 for PGA and 7 for the optimization method, while the optimal radial degree for both methods can be estimated to be 8 or 9.

**FIGURE 7.**
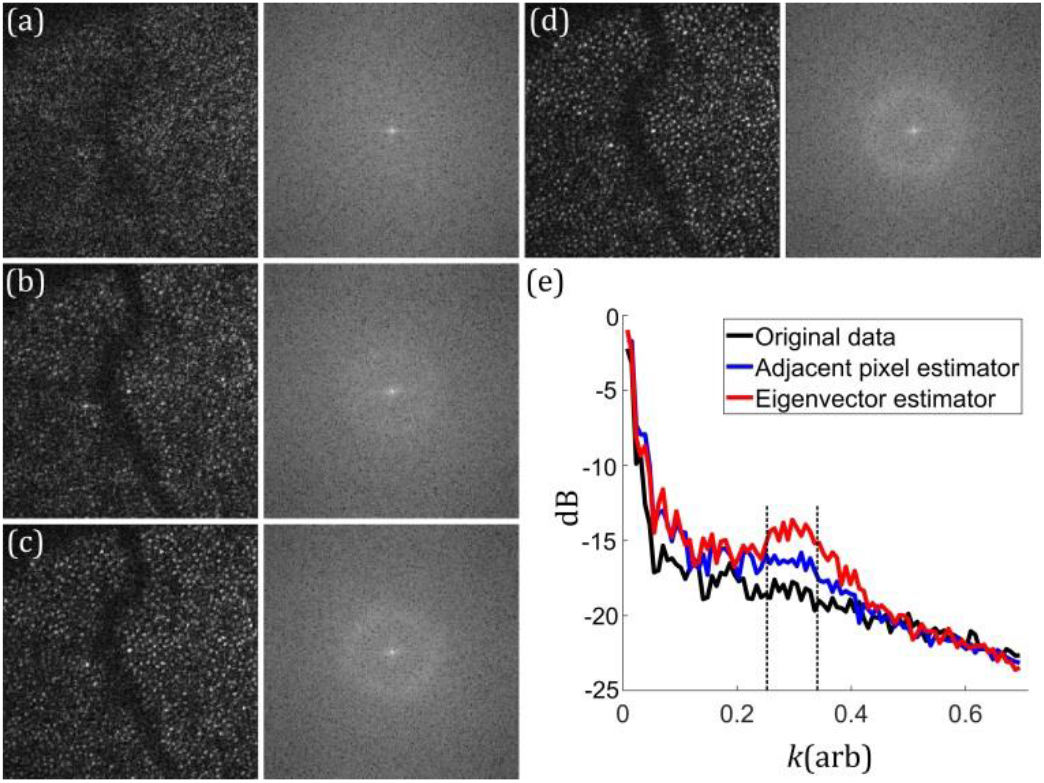
Comparing the performance of aberration correction using Yellott’s ring. a) Image and 2D Fourier transform before aberration correction. b)-d) 2D Fourier transform after aberration correction: b) with adjacent pixel estimator, c) with eigenvector estimator, d) with optimization method. e) black, blue, red lines the circle result of averaging around the centre of the Yellott’s rings corresponding to images a), b), c) respectively.

So, using polynomials up to radial degree 4 (12 polynomials, as in the work [14]) leads to a failure in the compensation procedure if applied to our data (Yellott’s ring is not visible with the use either of the proposed or of the optimization method). The application of the polynomials up to degree 6 (25 polynomials like in the work [15]), leads to a drop in the SNR by 1.1 dB with the use of the PGA method, while optimization method fails (apparently due to falling into a local minimum). In fig. 8 is presented the comparison of the algorithms performance with various maximum orders of Zernike polynomials

**FIGURE 8.**
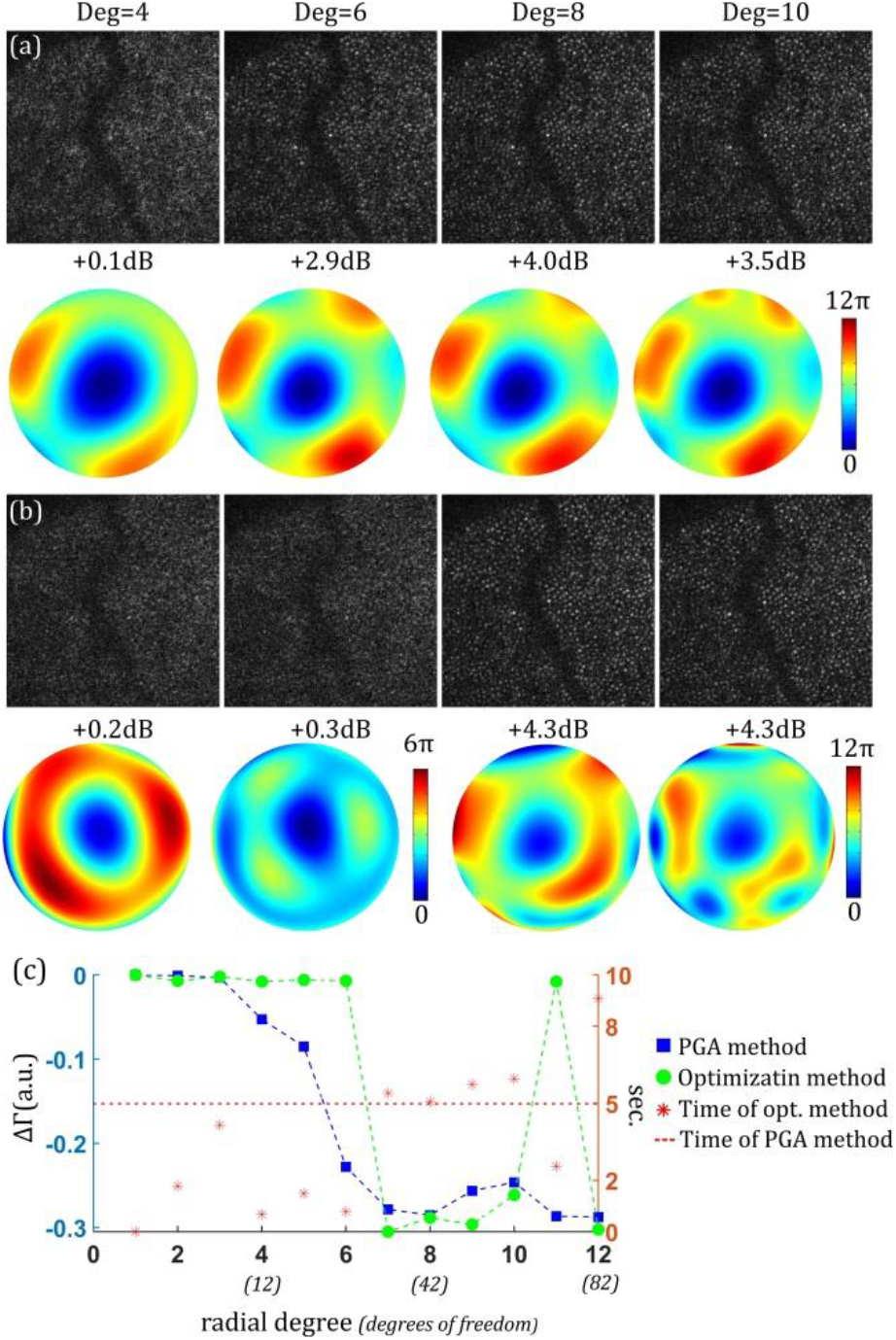
Comparison of the algorithms performance with various maximum orders of Zernike polynomials. Corrected images and determined aberrations are obtained: a) with PGA method b) with optimization method. The columns correspond to various maximum order (maximum degree of radial polynomial) of Zernike polynomials (4, 6, 8 and 10 respectively). Under the corrected images are presented SNR, calculated using Yellott’s ring (in comparison to the uncorrected image).c) Difference in Shannon entropy between the restored and initial images (left) and computational time (right) for the various radial degree of Zernike polynomials with correspondent number of the calculated coefficients (in parenthesis) (x-axis). Blue line with square markers is represented Shannon entropy for the PGA method, green line with circle markers represents Shannon entropy for the optimization method, red asterisks represent computational time for the optimization method, red dashed line represent computational time for the PGA method.

## 4 CONCLUSION

The efficiency of the PGA algorithm for compensating large eye aberrations has been demonstrated using real experimental data. The eigenvector estimator significantly increased the efficiency of the proposed method. The use of images with smaller, numerically reduced aperture proved to be critical for initial detection of sub-image centers. These two modifications, made numerical aberration correction for fundus imaging of the eye possible using PGA.

The algorithm ensures compensation comparable to that provided by the optimization method, but it works faster, especially for high-frequency wavefront distortions, which require a large number of optimization parameters. It seems reasonable to use PGA for detecting principal aberrations, and, if needed, to use the optimization method for refinement. In addition, to considerably increasing processing speed, it decreases the risk of finding a local minimum for the optimization method.

The advantage of the proposed method compared to the subaperture based methods is the ability to detect spatially high-frequency aberrations with a higher RMS deviation.

This method can be used on its own or in combination with the earlier proposed methods for improving the efficiency or image quality.

## ACKNOWLEDGMENTS

The research was supported by Contract of the Russian Federation RSF project no. 17-72-20249 and the German Research Foundation (HU 629/6-1).

